# Host Cell Proteases Drive Early or Late SARS-CoV-2 Penetration

**DOI:** 10.1101/2020.12.22.423906

**Authors:** Jana Koch, Zina M Uckeley, Patricio Doldan, Megan Stanifer, Steeve Boulant, Pierre-Yves Lozach

## Abstract

SARS-CoV-2 is a newly emerged coronavirus (CoV) that spread through human populations worldwide in early 2020. CoVs rely on host cell proteases for activation and infection. The trypsin-like protease TMPRSS2 at the cell surface, cathepsin L in endolysosomes, and furin in the Golgi have all been implicated in the SARS-CoV-2 proteolytic processing. Whether SARS-CoV-2 depends on endocytosis internalization and vacuolar acidification for infectious entry remains unclear. Here, we examined the dynamics of SARS-CoV-2 activation during the cell entry process in tissue culture. Using four cell lines representative of lung, colon, and kidney epithelial tissues, we found that TMPRSS2 determines the SARS-CoV-2 entry pathways. In TMPRSS2-positive cells, infection was sensitive to aprotinin, a TMPRSS2 inhibitor, but not to SB412515, a drug that impairs cathepsin L. Infectious penetration was marginally dependent on endosomal acidification, and the virus passed the protease-sensitive step within 10 min. In a marked contrast, in TMPRSS2-negative cells cathepsin L and low pH were required for SARS-CoV-2 entry. The cathepsin L-activated penetration occurred within 40-60 min after internalization and required intact endolysosomal functions. Importantly, pre-activation of the virus allowed it to bypass the need for endosomal acidification for viral fusion and productive entry. Overall, our results indicate that SARS-CoV-2 shares with other CoVs a strategy of differential use of host cell proteases for activation and infectious penetration. This study also highlights the importance of TMPRSS2 in dictating the entry pathway used by SARS-CoV-2.

**Significance:** Preventing SARS-CoV-2 spread requires approaches affecting early virus-host cell interactions before the virus enters and infects target cells. Host cell proteases are critical for coronavirus activation and infectious entry. Here, we reconcile apparent contradictory observations from recent reports on endosomal acidification and the role of furin, TMPRSS2, and cathepsin L in the productive entry and fusion process of SARS-CoV-2. Investigating authentic virus in various cell types, we demonstrated that SARS-CoV-2 developed the ability to use different entry pathways, depending on the proteases expressed by the target cell. Our results have strong implications for future research on the apparent broad tropism of the virus *in vivo*. This study also provides a handle to develop novel antiviral strategies aiming to block virus entry, as illustrated with the several drugs that we identified to prevent SARS-CoV-2 infection, some with low IC_50_.

## Introduction

The *Coronaviridae* is a large viral family of several hundred members, which constitutes along with *Arteriviridae* and *Roniviridae* the order *Nidovirales* (1). To date, four coronaviruses (CoVs) have been identified as the leading cause for common colds in humans (2). Three other CoVs, causing severe respiratory diseases, have emerged into the human population as a result of spillover events from wildlife during the last two decades (3). Severe acute respiratory syndrome (SARS)-CoV and Middle East respiratory syndrome (MERS)-CoV were first isolated in China in 2002 and Saudi Arabia in 2011, respectively (3). The most recent, SARS-CoV-2, is responsible for CoV induced disease (COVID-19) and turned into a pandemic in early 2020. As of December 22, 2020, more than 77 million human cases have been reported with at least 1.7 million deaths.

As other CoVs, SARS-CoV-2 particles are enveloped, roughly spherical, with a diameter between 90 and 110 nm (4, 5). The viral genome consists of one single-stranded positive-sense RNA segment that replicates in the cytosol and encodes four structural proteins. Three transmembrane proteins are embedded in the viral envelope and are exposed at the virion surface, namely the large glycoprotein S, the membrane protein M, and the envelope protein E (3). The nucleoprotein NP binds to the genomic RNA to form nucleocapsid structures inside the viral particles. In the viral envelope, glycoprotein S forms spike-like projections up to 35 nm in length, responsible for virus attachment to host cells and penetration by membrane fusion (6).

Although SARS-CoV-2 has been the subject of intense research since the beginning of 2020, our current understanding of cell entry remains essentially derived from studies on SARS-CoV and other CoVs (3). SARS-CoV-2 has been shown to rely on ACE2 (7), heparan sulfates (8), and neuropilin-1 (9) at the cell surface for infection. Inhibitor studies support the possibility that the virus enters the endosomal vesicles and relies on vacuolar acidification for the infectious entry process (7, 10, 11). As with many other CoVs, there is intense debate as to whether SARS-CoV-2 enters the host cells from the plasma membrane or from intracellular compartments.

To gain access into the cytosol, enveloped viruses must fuse their envelope with the cell membrane. Several classes of viral fusion proteins are known to mediate this process, each with their own molecular specificities [reviewed in (12)]. Structural studies categorized the SARS-CoV-2 protein S as a Class-I viral fusion protein, within the same group as other corona-, human immunodeficiency, and influenza (IAV) viruses (13–15). Cryo-electron microscopy showed that the S protein forms homotrimers at the surface of SARS-CoV-2 particles, in which the viral fusion subunits are buried (13, 14). The activation of the Class-I viral fusion proteins usually involves proteolytic processing, and membrane fusion is triggered by interactions with cell receptors and sometimes endosomal acidification. Activation and priming are irreversible steps, and the Class-I viral fusion proteins act only once (12). In the case of SARS-CoV-2, endosomal acidification appears to be non-essential to induce the spike-mediated fusion of the host membrane with the viral envelope (16). Yet why SARS-CoV-2 infection is sensitive to perturbants of endosomal acidification remains unclear.

Several proteases have been proposed to prime and activate the S protein (17), a step prior virus fusion and infection. Furin is a calcium-dependent serine endoprotease widely expressed in tissues. It has been proposed to cleave the S protein at the site S1/S2 (17–19), most likely when the viral progeny exits the infected cells. The cleavage results in two subunits, S1 and S2. S1 contains a receptor binding domain, and S2 the membrane fusion effector. An additional proteolytic cleavage in the S2 subunit occurs at the site S2’ during virus entry to trigger the fusion of the viral envelope with the host cell membrane. The transmembrane serine protease 2 (TMPRSS2), a cell surface trypsin-like protease (20), and cathepsin L, an endolysosomal cysteine protease (21), have both been proposed to be involved in the cleavage at the S2’ site (5, 7, 17, 22, 23). Still, the timing and dynamics of proteolytic cleavages and their potential role in SARS-CoV-2 activation, fusion, and entry remain unclear.

SARS-CoV-2 primarily targets cells of the lung epithelium but is also found in many other epithelial tissues as it spreads throughout the host. The fact that epithelia express ACE2, TMPRSS2, and cathepsin L most likely differentially influences the cell entry mechanisms of SARS-CoV-2 in a specific manner. In the present study, we developed sensitive, quantitative assays to analyze the SARS-CoV-2 entry process in different epithelial cell types. Using these assays, we determined SARS-CoV-2 dependence on low pH, proteolytic processing, proteases- and endosomal acidification-requiring dynamics, endocytosis, and protease-activated membrane fusion. Our work established that SARS-CoV-2 shares with MERS-CoV and other CoVs the ability to make a differential use of host cell proteases to enter and infect target cells.

## Results

### Characterization of SARS-CoV-2 Life Cycle in Caco-2 and Vero Cells

Many epithelial cell types have been reported to support productive SARS-CoV-2 infection (7), and both the TMPRSS2 and cathepsin L proteases have been implicated in the proteolytic processing of the viral S protein (5, 7, 17, 22, 23). We selected four epithelial cell lines that are known to support SARS-CoV-2 infection, i.e., Calu-3, Caco-2, A549, and Vero cells (7). A549 are intrinsically poorly infectable by SARS-CoV-2 due to the absence of the SARS-CoV-2 receptor ACE2 (7). As such, we used A549 cells stably overexpressing ACE2 (A549*). When cell lysates were subjected to SDS-PAGE and western blotting, we found that TMPRSS2 was effectively expressed in Calu-3 cells and to a lower extent in Caco-2 cells (Fig. 1*A*), corroborating results from others (24). Regardless of the presence of TMPRSS2, cathepsin L (from 25 to 31 kDa) and its inactive form, i.e., procathepsin L (35 to 41 kDa), were present in all the cell lines (Fig. 1*B*). However, the conversion of procathepsin L to cathepsin L appeared significantly higher in Vero cells than in the three other cell lines.

**Fig. 1.**
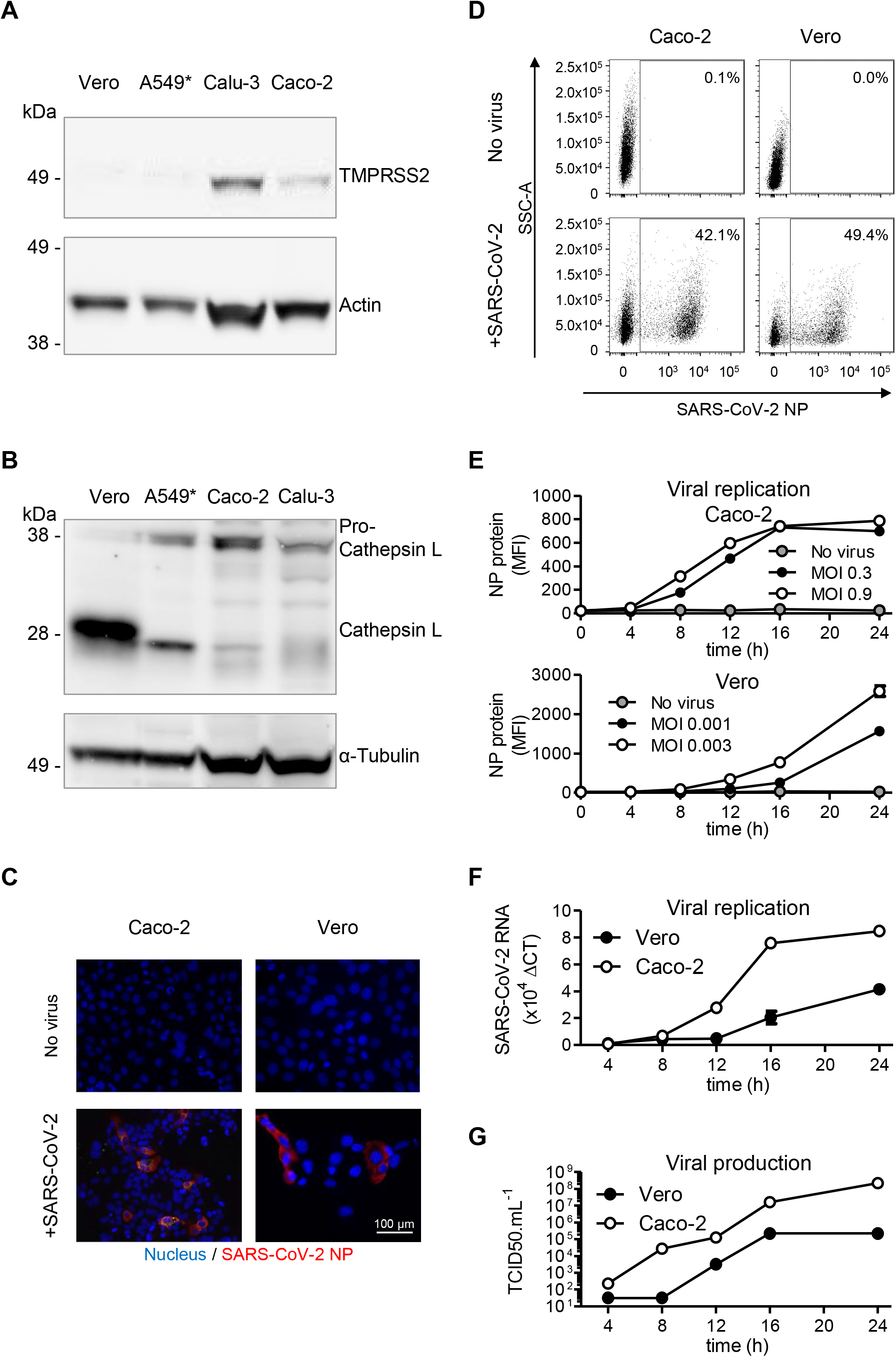
Quantification of SARS-CoV-2 infection. (*A* and *B*) Cells were lysed and analyzed by SDS-PAGE and western blotting under non-reducing conditions (A) and reducing conditions (B). A549*, ACE2-expressing A549 cells. (*C*) Vero and Caco-2 cells were infected with SARS-CoV-2 at a MOI of 0.5 and 0.1, respectively, for 8 h. Infected cells were then permeabilized and immunostained against the intracellular SARS-CoV-2 nucleoprotein (NP, red). Nuclei were stained with DAPI (blue) before imaging by fluorescence wide-field microscopy. (*D*) Vero and Caco-2 cells were exposed to SARS-CoV-2 at a MOI of 0.003 and 0.3, respectively, and harvested 16 h later. After fixation and permeabilization, infected cells were stained with the primary mAb against NP. Infection was analyzed by flow cytometry. SSC-A, side scatter, area. (*E*) Infection of Vero and Caco-2 cells was monitored over 24 h using the flow cytometry-based assay used for the experiment shown in panel D. Infection is given as the total fluorescence associated with the NP protein-positive cells. MFI, mean of fluorescence intensity. (*F*) SARS-CoV-2 mRNA levels were quantified by qRT-PCR in both Vero and Caco-2 cells infected at MOIs of 0.5 and 0.1, respectively, for up to 24 h. (*G*) Supernatants from infected cells were collected during the time course in F and assessed for the production of new infectious viral particles using a TCID50 assay on naïve Vero cells.

To address how the presence or absence of TMPRSS2 influences the SARS-CoV-2 infectious penetration, and how the endosomal acidification contributes to the process, we aimed to compare cell lines expressing or not this protease. To this end, we first defined the timing for a single round of infection using our cell lines. Calu-3 and Caco-2 served as TMPRSS2-positive (TMPRSS2+) cells, and A549* and Vero as non-expressing cells. The susceptibility of Caco-2 and Vero cells to SARS-CoV-2 at multiplicities of infection (MOIs) of 0.1 and 0.5, respectively, was assessed by fluorescence microscopy after immunostaining with a mouse monoclonal antibody (mAb) against the intracellular viral nucleoprotein NP (Fig. 1*C*). Results show that 10% of Caco-2 cells were positive for NP at 8 hours post-infection (hpi). Similarly, 35% of Vero cells were found infected at 8 hpi (Fig. 1*C*).

To quantify infection more accurately, we then performed flow cytometry analysis of Caco-2 and Vero cells infected with different MOIs of SARS-CoV-2 (Fig. 1*D* and *E*). The fluorescence increased over time and reached a plateau within 16 to 24 hpi (Fig. 1*E*), showing that the signal detected in the flow cytometry-based assays corresponded to viral replication and not to input particles. These kinetics were in agreement with real-time quantitative reverse transcription PCR (qRT-PCR) monitoring over time the amount of SARS-CoV-2 genome (Fig. 1*F*).

To evaluate the production and release of *de novo* infectious viral particles, we infected Caco-2 and Vero cells and quantified virus production up to 24 hpi by 50% tissue culture infective dose assay (TCID50). Infectious progeny viruses were found to be released from infected cells as early as 8-12 hpi (Fig. 1*G*). Virus replication kinetics and *de novo* virus release was found to be similar in Calu-3 and A549* cells (data not shown). Altogether, our analysis revealed that SARS-CoV-2 completes one round of infection, from virus binding and entry to replication and release of *de novo* infectious particles, within 8 h in Caco-2 cells and somewhat longer in Vero cells, i.e., between 8 and 12 h. In all the further experiments, as we aimed at characterizing SARS-CoV-2 entry mechanisms therefore we used MOIs allowing the infection of about 20% of the cells and limited our assays to 8 hpi.

### SARS-CoV-2 Makes a Differential Use of Host Cell Proteases for Infectious Penetration

To evaluate the role of the cell surface TMPRSS2 and endolysosomal cathepsin L proteases in the entry mechanisms of SARS-CoV-2, we used aprotinin and SB412515, respectively, to selectively inhibit the two proteases. As expected, no noticeable effect was observed when aprotinin was added to TMPRSS2-negative (TMPRSS2-) cells (A549* and Vero cells) prior to infection (Fig. 2*A*). In agreement with previous work (25), we observed that aprotinin reduced SARS-CoV-2 infection in a dose-dependent manner in the cells that express TMPRSS2 (Calu-3 and Caco-2 cells) (Fig. 2*A*). Conversely, SB412515 effectively prevented the infection of cells lacking TMPRSS2 (Vero and A549* cells) in a dose-dependent manner but had no effect on SARS-CoV-2 infection of Calu-3 and Caco-2 cells (Fig. 2*B*). The fact that aprotinin interfered with SARS-CoV-2 infection in Calu-3 and Caco-2 cells indicated that, even if TMPRSS2 was blocked, cathepsin L would not take over and subsequently process SARS-CoV-2.

**Fig. 2.**
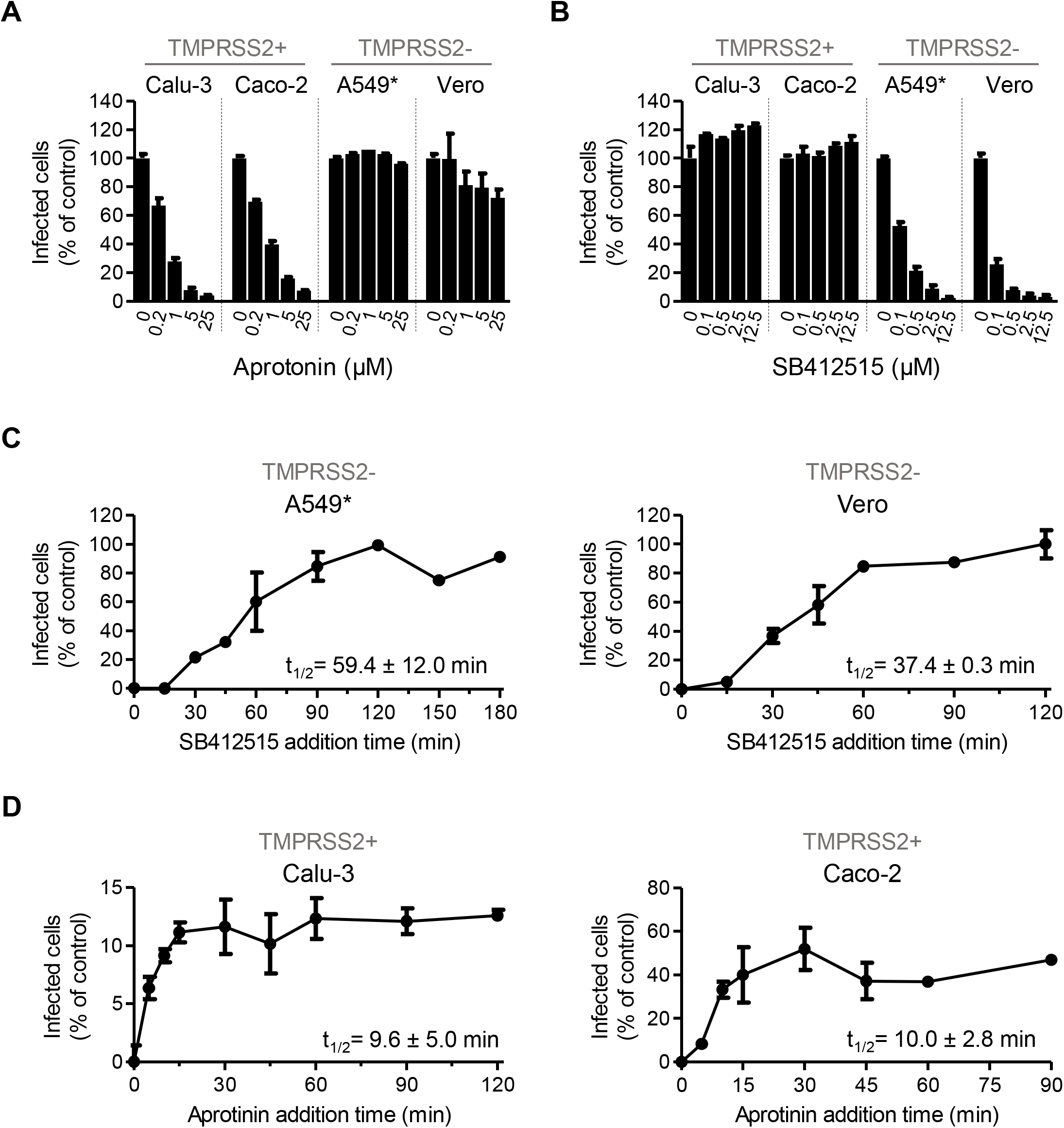
SARS-CoV-2 makes a differential use of host cell proteases for infectious penetration. (*A* and *B*) Cells were pre-treated at indicated concentrations of aprotinin (A) and SB412515 (B), which are inhibitors of TMPRSS2 and cathepsin L, respectively. Infection with SARS-CoV-2 (MOI of 0.9) was achieved in the continuous presence of drug. Infected cells were quantified by flow cytometry as described in Fig. 1*D*, and data normalized to samples where inhibitors had been omitted. (*C* and *D*) SARS-CoV-2 particles (MOI of 0.9) were bound to A549* and Vero cells (C) or Calu-3 and Caco-2 cells (D) on ice for 90 min, and subsequently, warmed rapidly to 37°C to allow infectious penetration. 10 μM of SB412515 (C) or 30 μM of aprotinin (D) were added at different times post warming to block further proteolytic activation. Infection was analyzed by flow cytometry, and data were normalized to samples where protease inhibitors had been omitted.

We next determined the kinetics of the cathepsin L- and TMPRSS2-dependent SARS-CoV-2 entry process. Cells were incubated with viruses at a low MOI (~0.9) on ice and rapidly shifted to 37°C to allow virus entry and protease activity. The cathepsin L and TMPRSS2 inhibitors were added at different times after warming to prevent further activation and penetration of the virus. In other words, we determined the time when inhibition of SARS-CoV-2 activation is no longer possible, which resulted in an increase of infection. In both TMPRSS2-cell lines (A549* and Vero cells), the SB412515 add-in time course revealed that the activation by cathepsin L and the subsequent infectious penetration of SARS-CoV-2 started after a 15 min lag and reached a half maximal level (t_1/2_) within 40-60 min (Fig. 2*C*). Evidently, exposure of individual viruses to cathepsin L occurred non-synchronously during a time span 15-90 min after warming. The add-in time course using aprotinin showed that productive penetration was much faster in TMPRSS2+ cells (Calu-3 and Caco-2 cells) (Fig. 2*D*). The t_1/2_ of activation by TMPRSS2 was reached within 5-10 min in both cell lines. Taken together, our observations demonstrated that TMPRSS2 allowed for a faster activation and penetration of SARS-CoV-2 in comparison to cells for which infection depends on cathepsin L.

### TMPRSS2 Governs SARS-CoV-2 Dependence on Low pH for Infectious Entry

Recent reports indicated that SARS-CoV-2 infection is sensitive to lysosomotropic weak bases that neutralize vacuolar pH such as ammonium chloride (NH4Cl) and chloroquine (7, 10, 11). However, TMPRSS2 is active at the cell surface under neutral pH conditions (20), unlike cathepsin L, which requires the low-pH environment typical of endolysosomes (21). To assess the importance of endosomal acidification for infectious entry in cells expressing (Caco-2 and Calu-3) and lacking (A549* and Vero) TMPRSS2, cells were exposed to SARS-CoV-2 in the presence of increasing amounts of NH_4_Cl or chloroquine. Our results showed that both weak bases induced a dose-dependent inhibition of infection regardless of the cell type and of TMPRSS2 expression (Fig. 3*A* and 3*B*). However, the dose to inhibit 50% of SARS-CoV-2 infection (IC_50_) was found to be significantly lower in cells devoid of TMPRSS2 compared to cells expressing the protease, reaching a 200-fold difference for chloroquine (Table 1).

**Fig. 3.**
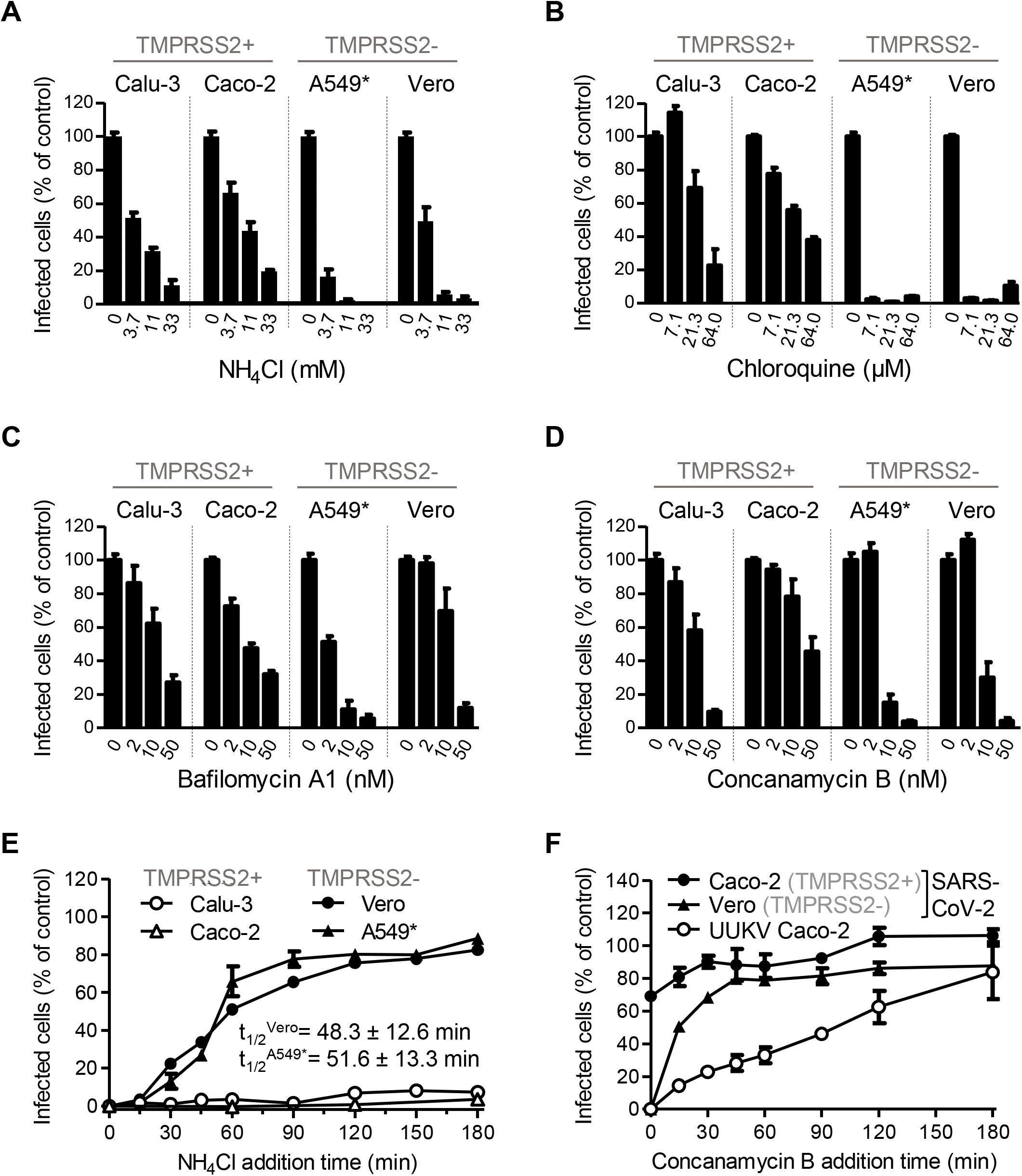
SARS-CoV-2 infection depends on endosomal acidification. (*A* to *D*) Cells were pre-treated with endosomal-pH interfering drugs at indicated concentrations and subsequently infected with SARS-CoV-2 in the continuous presence of drug, namely NH_4_Cl (A), chloroquine (B), Bafilomycin A1 (C), Concanamycin B (D). Infected cells were quantified by flow cytometry as described in Fig. 1*D*, and data normalized to samples where inhibitors had been omitted. (*E*) Binding of SARS-CoV-2 to cells was synchronized on ice for 90 min. Subsequently, cells were rapidly shifted to 37°C to allow penetration. NH_4_Cl (50 mM for A549* and Vero cells, and 75 mM for Calu-3 and Caco-2 cells) was added at indicated times to neutralize endosomal pH and block the acid-dependent step of SARS-CoV-2 infectious penetration. Infected cells were analyzed by flow cytometry, and data normalized to samples where NH_4_Cl had been omitted. (*F*) Same than in (E) but using Concanamycin B (50 nM) instead NH_4_Cl. Uukuniemi virus (UUKV) was used to control the efficiency of Concanamycin B to neutralize endosomal pH in Caco-2 cells.

**Table 1.** Half maximal inhibitory (IC_50_) of inhibitors against SARS-CoV-2.

|  | **Calu-3** | **Caco-2** | **A549\*** | **Vero** |
| --- | --- | --- | --- | --- |
| Aprotonin | 0.4 ± 0.1 µM | 0.6 ± 0.0 µM | x | x |
| SB412515 | x | x | 125.7 ± 29.9 nM | 36.9 ± 10.9 nM |
| NH<sub>4</sub>Cl | 4.4 ± 0.9 mM | 7.9 ± 2.4 mM | 2.2 ± 0.1 mM | 2.5 ± 0.6 mM |
| Chloroquine | 50.1 ± 24.4 µM | 27.4 ± 4.0 µM | 0.3 ± 0.0 µM | 0.2 ± 0.1 µM |
| Bafilomycin A1 | 16.3 ± 6.6 nM | 10.4 ± 3.2 nM | 2.0 ± 0.6 nM | 18.6 ± 7.7 nM |
| Concanamycin B | 12.2 ± 5.8 nM | 50.3 ± 30.4 nM | 6.0 ± 1.2 nM | 8.6 ± 2.2 nM |
| MG-132 | 0.7 ± 0.2 µM | 5.2 ± 2.1 µM | 4.4 ± 1.4 nM | 16.4 ± 5.6 nM |

To validate the observation that TMPRSS2+ cells were less dependent on endosomal acidification for SARS-CoV-2 infection, we made used of Bafilomycin A1 and Concanamycin B, which are inhibitors of the vacuolar-type proton-ATPases (vATPases). Incubation of cells with increasing amounts of the two drugs resulted in a dose-dependent inhibition of SARS-CoV-2 infection (Fig. 3*C* and 3*D*). Importantly, the inhibition was marginal with 10 nM of Bafilomycin A1 and Concanamycin B in TMPRSS2+ cells (Caco-2 and Calu-3), and the decrease in infection did not exceed 50-80% at 50 nM of Concanamycin B. For comparison, infection with Uukuniemi virus (UUKV), a late-penetrating virus that relies on low pH in late endosomes (LE) for penetration (26), is strongly inhibited in the presence of 2 to 10 nM of Concanamycin B or Bafilomycin A1 (26). From these results, it was evident that, similar to the lysosomotropic weak bases, SARS-CoV-2 infection appeared to be significantly less sensitive to vATPase inhibitors in TMPRSS2+ cells (Caco-2 and Calu-3) in comparison to cells lacking the protease (Vero and A549* cells) (Table 1).

### SARS-CoV-2 Can Use Two Distinct Routes to Enter and Infect Target Cells

Our results suggested that SARS-CoV-2 infection relied more on endosomal acidification in cells devoid of TMPRSS2 than cells expressing the protease. To pursue this possibility, we determined the kinetics of the acidification step required for the infectious penetration of SARS-CoV-2 into TMPRSS2-cells. We took advantage of the fact that the neutralization of endosomal pH is nearly instantaneous upon NH_4_Cl addition to the extracellular medium (27). Virus particles were first allowed to attach to A549* and Vero cells on ice. Entry was then synchronized by switching cells rapidly to 37°C, and NH_4_Cl was added at different times. In A549* and Vero cells, viruses passed the NH_4_Cl-sensitive step 15 min after cell warming, and the t_1/2_ was reached within 50 min (Fig. 3*E*). Overall, the kinetics of SARS-CoV-2 acid-activated penetration closely resembled the time course of cathepsin L-dependent activation in the absence of TMPRSS2 (Fig. 2*C*).

In Calu-3 and Caco-2 cells, both of which express TMPRSS2, it was not possible to determine the timing of the acid-requiring step. We failed to detect SARS-CoV-2-infected cells even by adding NH_4_Cl several hours after transferring the cells from 4 to 37°C (Fig. 3*E*). In samples where NH_4_Cl was omitted, infection was readily detectable with 17% of Calu-3 and Caco-2 cells infected (data not shown) suggesting that the weak base interferes with SARS-CoV-2 replication in these two cell lines. It is highly likely that NH_4_Cl disrupts TMPRSS2+ cell-specific functions that are important for SARS-CoV-2 replication. NH_4_Cl not only neutralizes the intracellular pH but also alters all endosomal, lysosomal, and trans-Golgi-network functions that are acid dependent (28).

As an alternative method to alter endosomal pH, we used Concanamycin B instead of NH_4_Cl and added the vATPase inhibitor to Caco-2-cell-bound virus at different times after warming. The time course showed that SARS-CoV-2 infection was insensitive to the Concanamycin B add-in as early as a few seconds after shifting Caco-2 cells to 37°C (Fig. 3*F*). In a marked contrast, infectious entry of UUKV started after 15 min and had not reached a maximum 2 h after cell warming. As expected, SARS-CoV-2 passed the Concanamycin B-sensitive step in Vero cells within less than 15 min, and infectious entry reached a plateau value after 45 min, somewhat faster than in using NH_4_Cl (Fig. 3*E*). This difference in Vero sensitivity to endosomal pH may be that Concanamycin B not only interferes with endosomal functions that are acid dependent but also indirectly with the maturation of endosomes. However, unlike NH_4_Cl, it was apparent that Concanamycin B had no adverse effect on SARS-CoV-2 replication in all these experiments. Taken together, these results strongly suggested that SARS-CoV-2 can use two different routes to enter and infect target cells, i.e., fast pH-independent penetration in TMPRSS2+ cells (Fig. 2*D* and 3*F*) and slow acid-activated entry in cells lacking TMPRSS2 (Fig. 2*C* and 3*E*).

### SARS-CoV-2 Relies on Endolysosomal Maturation for Infection of TMPRSS2-Cells

The timing of acid-dependent and protease-activated steps suggested that SARS-CoV-2 penetration might occur from endolysosomes in cells devoid of TMPRSS2 and from the plasma membrane or early endosomes (EEs) in TMPRSS2+ cells. To determine whether SARS-CoV-2 requires reaching the endolysosomal compartments for the productive infection of TMPRSS2-cells, we exploited the small GTPase Rab7a, which is a key player of LE maturation and function. TMPRSS2-Vero cells were transfected with DNA plasmids encoding the wild-type (wt), the dominant-negative (Rab7a T22N), and the constitutively active (Rab7a Q67L) forms of Rab7a tagged with the enhanced green fluorescent protein (EGFP) prior to infection with SARS-CoV-2. Transfected cells were selected for different levels of EGFP expression and then analyzed for infection. Increasing expression of the wt molecule of Rab7a facilitated SARS-CoV-2 infection. On the contrary, increasing expression of both mutants of Rab7a, which abrogates the maturation of newly formed LEs (26, 29), resulted in a 50% decrease in infection (Fig. 4*A*), indicating that the virus cannot fuse in Rab7a T22N- and Q67L-late endosomal vesicles. This result suggested that proper maturation of LEs is mandatory for the cathepsin L-dependent infectious entry of SARS-CoV-2.

**Fig. 4.**
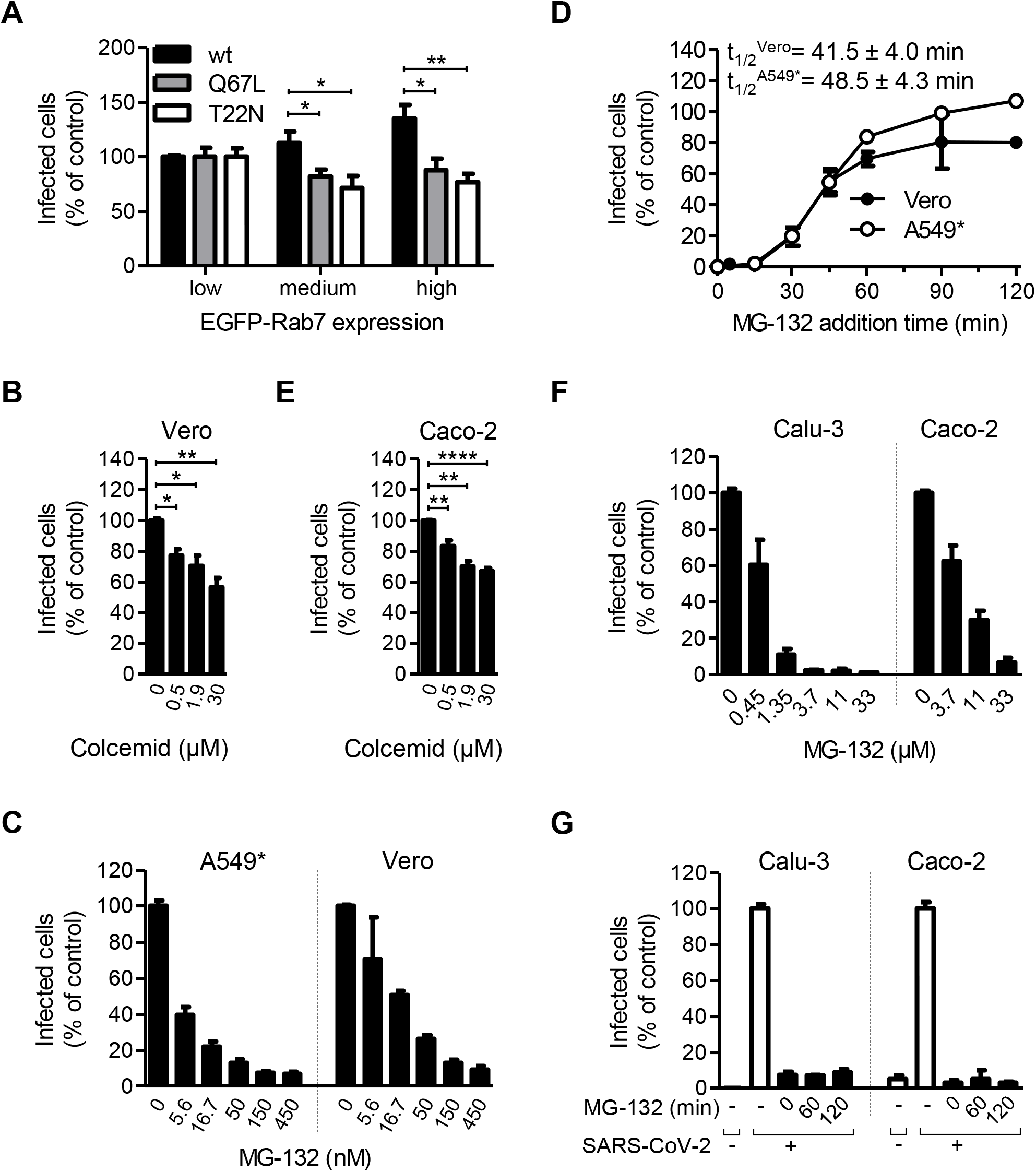
SARS-CoV-2 relies on late endosomal maturation for infection. (*A*) EGFP-Rab7a wild-type (wt), Q79L (constitutively active mutant), and T22N (dominant-negative mutant) were transiently expressed in Vero cells. The cells were then infected with SARS-CoV-2 at a MOI ~0.003. Using flow cytometry, cell populations were selected for levels of EGFP-Rab7a expression in roughly one-log increments, and infected cells were quantified within each population 8 hpi. Data were normalized to infection in cell populations with the lowest EGFP-Rab7a intensity. Unpaired t-test with Welch’s correction was applied. *, p < 0.05; **, p < 0.01. RU, relative unit. (*B* and *C*) Cells were pre-treated with colcemid (*B*) and MG-132 (*C*) at indicated concentrations and subsequently infected with SARS-CoV-2 in the continuous presence of inhibitors. Infection was analyzed by flow cytometry, and data were normalized to samples where inhibitors had been omitted. Unpaired t-test with Welch’s correction was applied. *, p < 0.05; **, p < 0.01; ****, p < 0.0001. (*D*) SARS-CoV-2 particles (MOI of 0.9) were bound to A549* and Vero cells on ice for 90 min, and then, switched rapidly to 37°C to allow infectious penetration. MG-132 (3.7 μM) was added to cells at indicated times to block further late endosomal maturation. Infection was analyzed by flow cytometry, and data were normalized to samples where MG-132 had been omitted. (*E*) As in the panel B but using Caco-2 cells instead Vero cells. (*F*) Same as C, except for Calu-3 and Caco-2 cells. (*G*) The timing of the MG-132-sensitive step during SARS-CoV-2 infectious entry into Calu-3 and Caco-2 cells was assayed as detailed in D but using 60 μM of MG-132.

LE maturation relies on microtubule-mediated transport to the nuclear periphery and proteasome activity (26, 29). Treatment of Vero cells with colcemid, a drug that interferes with microtubule polymerization, resulted in a 30%-45% decrease in infection (Fig. 4*B*). Additionally, late endosomal penetration of IAV and UUKV has been shown to be sensitive to free ubiquitin depletion produced by the proteasome inhibitor MG-132 (26, 30). Therefore, to determine if free ubiquitin was required for SARS-CoV-2 infection, A549* and Vero cells were treated with MG-132. Results show that SARS-CoV-2 infection was strongly inhibited in the presence of MG-132 in both cell lines (Fig. 4*C*). The calculated IC_50_ confirmed the high proficiency (4 to 17 nM) of MG-132 to interfere with the cathepsin L-mediated SARS-CoV-2 entry route (Table 1).

To determine the kinetic of the MG-132-sensitive step in the entry process, we followed the same experimental procedure used to determine the kinetics of endosomal acidification-dependent and cathepsin L-mediated activation of SARS-CoV-2 (Fig. 2*C* and 3*E*) but utilizing MG-132 instead of protease inhibitor and NH_4_Cl. Briefly, viruses were bound to A549* and Vero cells at a low MOI on ice, and then promptly switched to 37°C before adding MG-132 at different times. After a 15 min lag, infectious penetration occurred asynchronously between 30 and 60 min, with a t1/2 within 40-50 min (Fig. 4*D*). This time course was consistent with endolysosomal maturation, which usually lasts 30-60 min (31). Altogether, these results show that the cathepsin L-dependent SARS-CoV-2 infection depends on endolysosome maturation in TMPRSS2-A549* and Vero cells.

Interestingly, LE maturation was also required in TMPRSS2+ cells (Calu-3 and Caco-2 cells) as SARS-CoV-2 infection was reduced by colcemid in a dose-dependent manner in both cell lines (Fig. 4*E*). Though the inhibition was efficient, the IC_50_ values of MG-132 were one to three logs higher in TMPRSS2+ cells compared to TMPRSS2-cells (Fig. 4*F*). As shown in Fig. 4*G*, infection of Calu-3 and Caco-2 cells was not readily detectable when MG-132 was added 2 hpi. Together, the data suggest that MG-132 impaired viral replication in these assays and not the TMPRSS2-dependent SARS-CoV-2 entry process.

### Low pH Is Not Essential for SARS-CoV-2 Membrane Fusion

The penetration of enveloped viruses into the cytosol involves fusion between the viral envelope and a cell membrane. In most cases, endosomal acidification contributes to activate viral glycoproteins and is used as a cue to trigger fusion (12). SARS-CoV-2 does not rely on endosomal acidification to enter TMPRSS2+ cells, which suggests that the virus does not rely on low pH for membrane fusion but solely for the activation of cathepsin L in cells lacking TMPRSS2.

To examine the requirement of pH acidification for the SARS-CoV-2 membrane fusion mechanisms, we first assessed the possibility to inactivate the virus with acidic buffers prior infection. In such an assay, the virus undergoes a transition toward the post-fusion state at the optimal pH. If the transition is irreversible, the spike protein is no longer able to fuse with target-cell membranes, and thus, the viral particles are rendered non-infectious. With this approach, we found that about 50% of viruses were still infectious in Caco-2 and Vero cells, even after an exposition to buffers at pH ~5 for 10 min (Fig. 5*A*). Semliki forest virus (SFV) is an early-penetrating virus that has a Class-II viral fusion glycoprotein with an irreversible priming step triggered at a pH-activation threshold of 6.2 (26). In contrast to SARS-CoV-2, infection by low pH-pretreated SFV was reduced by 70-80% at pH ~6.0 and below (Fig. 5*B*).

**Fig. 5.**
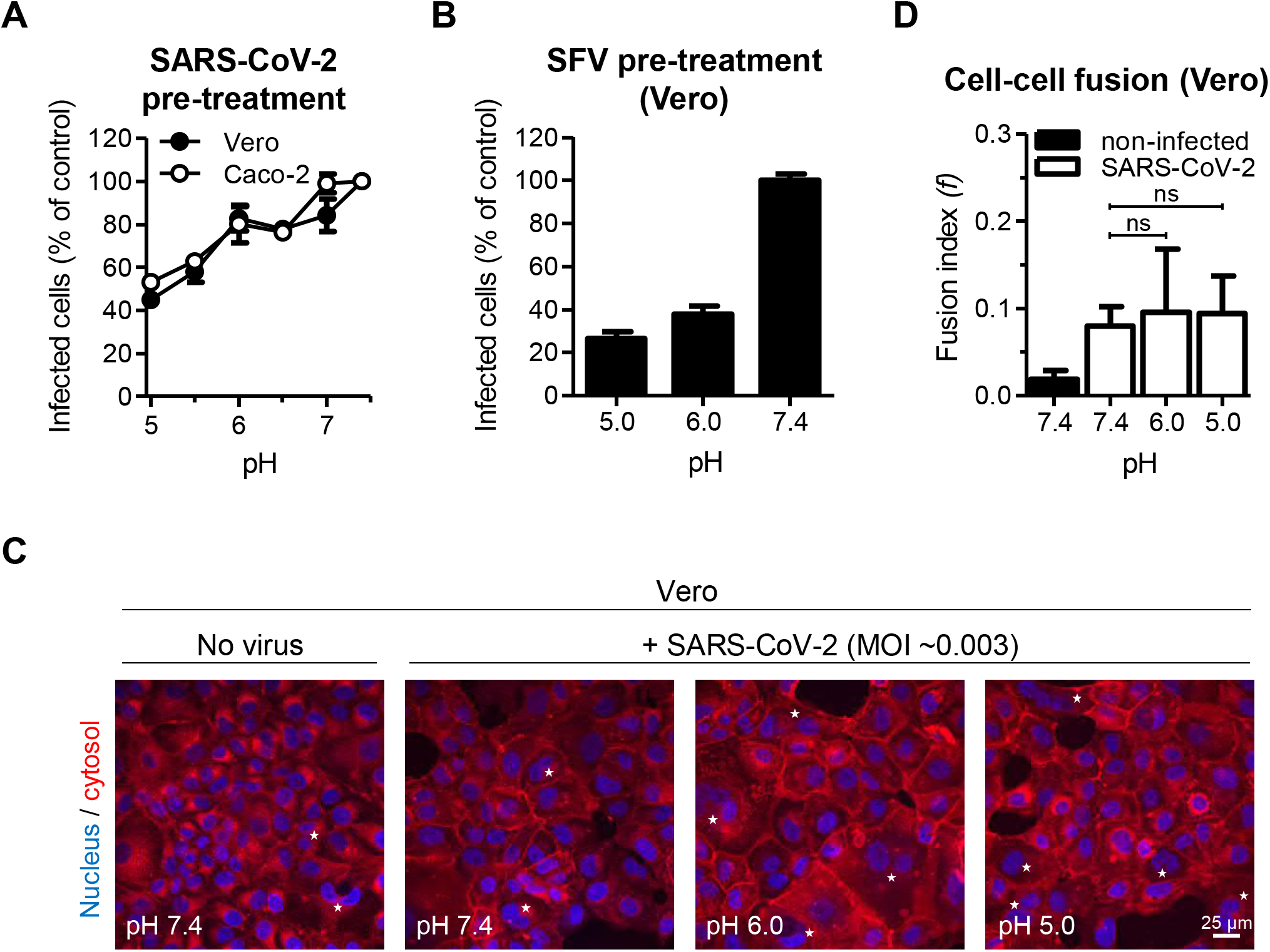
Acidification is not sufficient to trigger SARS-CoV-2 membrane fusion. (*A*) SARS-CoV-2 and (*B*) Semliki forest virus (SFV) particles were pre-treated at indicated pH for 10 min at 37°C. Viruses were subsequently neutralized with buffers at pH ~7.4 and allowed to infect Caco-2 and Vero cells. Infected cells were then immunostained against the NP protein and analyzed by flow cytometry. Data are normalized to samples pretreated with buffers at pH ~7.4. (*C*) Confluent monolayers of Vero cells were infected with SARS-CoV-2 at a MOI ~0.003 for 24 h prior to treatment with buffers at indicated pH for 5 min at 37°C. Plasma membrane was stained 1 h post-treatment with CellMask Deep Red (red). After fixation, nuclei were stained with Hoechst (blue). White stars indicate syncytia. (*D*) Images of microscope fields (32 < n < 44) obtained in (C) were quantified. Fusion index is given as *f* = 1 – [(number of cells in a field after fusion]/[number of nuclei)]. Unpaired t-test with Welch’s correction was applied. ns, non-significant.

To further investigate the influence of low pH on SARS-CoV-2 fusion, we then evaluated the capacity of SARS-CoV-2 to mediate cell-cell fusion (‘‘fusion-from-within’’) as described for unrelated viruses (32). To this end, we used Vero cells as they are negative for TMPRSS2, which makes it a convenient model to monitor proteolytic activation of the S protein at the cell surface by exogenous proteases. Briefly, confluent monolayers of Vero cells were infected with SARS-CoV-2 for 24 h, and the infected cells were then subjected to buffers of different pH values. The extent of cell-cell fusion, i.e., formation of syncytia, was determined using a fusion index that expresses the average number of fusion events per original mononucleated cell (33). The index reaches 1 when all the nuclei in the microscope field are present in a single cell, and the value is 0 when all cells have one nucleus each. Formation of syncytia with two or more nuclei was observed regardless of the pH of the buffer (Fig. 5*C*), and the fusion index did not significantly differ when cells were treated with low pH or neutral buffers (Fig. 5*D*). Together, our observations strongly suggested that low pH is not required for the SARS-CoV-2 fusion mechanisms.

### Proteolytic Processing Is Sufficient and Necessary for SARS-CoV-2 Fusion

The results suggested that acidification is not required to prompt viral fusion and that proteolytic processing might be sufficient. As furin and TMPRSS2 are believed to mediate the activation of the SARS-CoV-2 spike proteins, we then evaluated the ability of the two proteases to trigger SARS-CoV-2 activation and fusion using our flow cytometry-based infection analysis and syncytia-forming assay. In the following series of experiments, exogenous trypsin was used to mimic TMPRSS2 at the cell surface as the two enzymes are closely related and both belong to the group of trypsin-like proteases. The use of exogenous cathepsin L was excluded because the enzyme is only active at pH ~5, which would have made it impossible to distinguish between an effect due to low pH or proteolytic cleavage.

Viral particles were first subjected to proteases prior to being added to Caco-2 and Vero cells. We found that infection increased as much as 2- to 3-fold following the SARS-CoV-2 proteolytic processing by trypsin, whereas the pre-exposure of particles to furin had no apparent effect (Fig. 6*A*). Similar results were obtained with our cell-cell fusion assay. Large syncytia with five or more nuclei were observed when infected Vero cells were exposed to trypsin (Fig. 6*B*). Contrary to trypsin-treated cells, no difference was observed after furin treatment in comparison to the mock-treated samples, for which the only cells with more than one nucleus were those dividing (Fig. 6*B*). Additionally, the fusion index in Vero cells was increased under trypsin treatment compared to mock- and furin-treated cells (Fig. 6*C*). Altogether our data indicated that proteolytic cleavage is sufficient and necessary for SARS-CoV-2 membrane fusion.

**Fig. 6.**
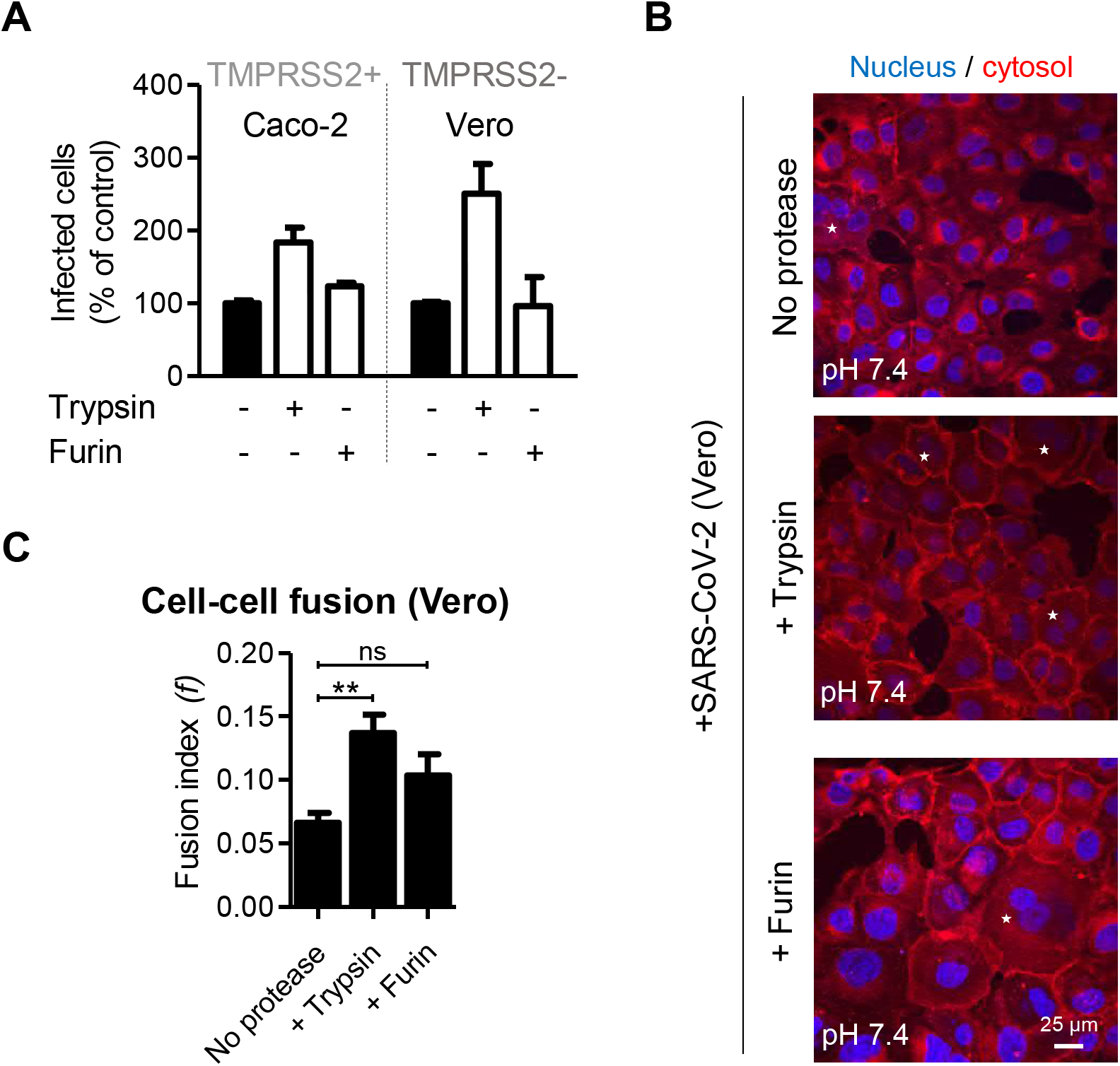
Proteolytic processing triggers SARS-CoV-2 membrane fusion. (*A*) SARS-CoV-2 (MOI of 1.2) was subjected to pretreatment with trypsin and furin for 15 min at 37°C prior infection of Caco-2 and Vero cells. Infected cells were quantified by flow cytometry as described in Fig. 1*D*. Data were normalized to samples not pre-treated with trypsin. (*B*) Confluent monolayers of Vero cells were infected with SARS-CoV-2 at a MOI ~0.003 for 24 h prior trypsin and furin treatment for 5 min at 37°C. Plasma membrane was stained with CellMask Deep Red (red) 1 h after trypsinization. After fixation, nuclei were stained with Hoechst (blue), and cells imaged by wide-field fluorescence microscopy. White stars indicate syncytia. (*C*) Images of microscope fields (n = 39, no protease, n = 63, +trypsin, and n = 54, +furin) obtained in (B) were quantified. Fusion index is calculated as in Fig. 5*D*. Unpaired t-tests with Welch’s correction was applied. **, p < 0.01.

### Endosomal Acidification Is Required for Endolysosomal Proteases Priming Viral Fusion

Our results support a model where endosomal acidification is not essential for SARS-CoV-2 membrane fusion, but SARS-CoV-2 infection relies on low pH for cathepsin L-dependent infection in cells lacking TMPRSS2. Therefore, we tested the possibility that acid pH is required for the activation of endolysosomal proteases that in turn trigger SARS-CoV-2 fusion. In such a scenario, the spike S proteins that are already primed by proteases should no longer rely on low pH for fusion. Indeed, we found that the fusion index was not increased when trypsin treatment was followed by exposure to a decreasing pH of 7.4 to 5 (Fig. 7*A* and 7*B*), the latter value being typical of the luminal pH of endolysosomes (29).

**Fig. 7.**
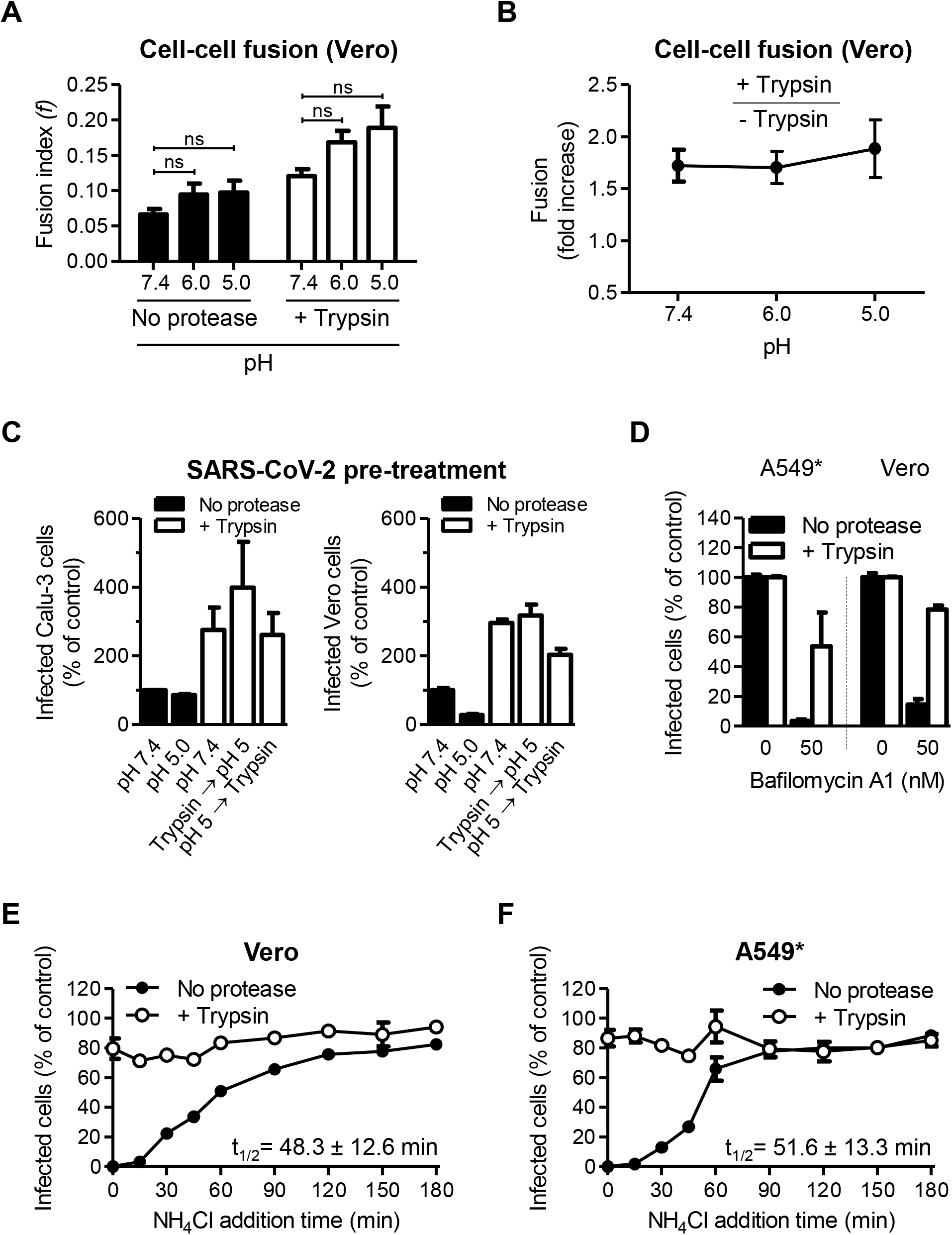
SARS-CoV-2 no longer requires endosomal acidification after proteolytic processing. (*A*) Confluent monolayers of Vero cells were infected with SARS-CoV-2 at a MOI ~0.003 for 24 h and then subjected to trypsin treatment for 5 min at 37°C. The cells were allowed to recover for 1 h at 37°C, and subsequently, exposed to buffers at indicated pH for 5 min at 37°C. Cell-cell fusion was determined as described in Fig. 6*B* and 6*C*. n > 28 microscope fields were analyzed, and unpaired t-test with Welch’s correction was applied. ns, non-significant. (*B*) Shows the increase in cell-cell fusion after trypsin treatment according to pH. The fusion is given as the ratio between the values obtained for trypsin-treated samples and those obtained for untreated samples. (*C*) SARS-CoV-2 particles (MOI of 1.2) were first subjected to trypsin treatment for 15 min at 37°C followed by exposition to buffers at indicated pH for 10 min at 37°C, and *vice versa*. A549* and Vero cells were then infected and analyzed by flow cytometry as described in Fig. 1*D*. (*D*) Trypsin-activated SARS-CoV-2 (MOI ~0.003) was allowed to infect A549* and Vero cells in the continuous presence of Bafilomycin A1. Infection was quantified by flow cytometry, and data normalized to samples where the inhibitor had been omitted. (*E* and *F*) Binding of trypsin-activated SARS-CoV-2 (MOI ~0.003) to Vero (E) and A549* (F) was synchronized on ice for 90 min. Subsequently, cells were rapidly shifted to 37°C to allow penetration. NH_4_Cl (50 mM) was added at indicated time to neutralize endosomal pH and block the acid-dependent step of SARS-CoV-2 infectious penetration. Infected cells were analyzed by flow cytometry, and data normalized to samples where NH_4_Cl had been omitted.

To further evaluate whether SARS-CoV-2 membrane fusion is low pH-independent, viral particles were then exposed to buffers at pH ~5 and subsequently subjected to proteolytic cleavage by trypsin. Our results revealed that SARS-CoV-2 infectivity was preserved when viral particles were exposed to the low-pH buffer prior to trypsin treatment in comparison to virus particles that were solely exposed to acidic pH (Fig. 7*C*). The infectivity also remained preserved when the virus was first subjected to trypsin and then acidification. Taken together, the results showed that endosomal acidification does not play a role in SARS-CoV-2 membrane fusion, whether it occurs before or after the proteolytic processing of viral particles. In addition, our results strongly suggested that the potential pH-induced conformational changes in the SARS-CoV-2 spikes were neither irreversible nor detrimental for the viral fusion.

To directly test whether endosomal acidification is needed for the host cell proteases that prime SARS-CoV-2 fusion, and not for the fusion mechanisms themselves, we assessed whether preactivated viral particles no longer depend on endosomal acidification for infectious entry. For this purpose, the proteolytic processing of the virus particles was achieved with trypsin prior to the infection of A549* and Vero cells. To interfere with the acid-dependent endolysosomal proteases, the infection was carried out in the continuous presence of 50 nM of Bafilomycin A1. As A549* and Vero cells do not express TMPRSS2, this assay allowed us to directly test the impact of extracellular protease-activated viral particles. As reported above (Fig. 3*C*), infection with untreated viral particles was severely hampered when proton pumps were blocked in the absence of TMPRSS2 (Fig. 7*D*). In stark contrast, the protease-preactivated viral particles remained infectious in the presence of Bafilomycin A1 (Fig. 7*D*).

The capacity of SARS-CoV-2 to infect A549* and Vero cells upon proteolytic activation, despite the absence of functional endolysosomal proteases, was confirmed using NH_4_Cl. As expected, in our synchronized infection assay, untreated particles became NH_4_Cl insensitive 50 min post entry (Fig. 7*E*, 7*F*, and 3*E*). However, when the viral particles were pretreated with trypsin, no sensitivity to NH_4_Cl was observed (Fig. 7*E* and 7*F*). These results strongly supported the view that, once activated by proteolytic cleavage, the virus is no longer dependent on endosomal acidification for infection. Altogether, our data show that SARS-CoV-2 resembles other CoVs in that its entry depends on diverse host cell proteases. It can use two distinct routes, where either TMPRSS2 mediates its pH-independent penetration from or close to the cell surface or alternatively, it is transported to endolysosomes, where low pH activates cathepsin L that in turn primes viral fusion and penetration.

## Discussion

The infectious entry process of CoVs is complex (3). Several host cell proteases can prime the CoV spike S proteins for viral membrane fusion, but it is not yet known whether these mechanisms require selective proteases or a coordinated, spatio-temporal combination of several proteases. The importance of endosomal acidification in the productive penetration of all CoVs is also a matter of debate. Furin, TMPRSS2, and cathepsin L have all three been implicated in coronavirus activation for entry (5, 7, 17, 22, 23), and agents elevating endosomal pH such as chloroquine have been described to interfere with infection (7, 10, 11). SARS-CoV-2 and other CoVs have apparently found a way to use diverse entry mechanisms to infect target cells and spread throughout the host.

In this study, we developed reliable and accurate assays to investigate SARS-CoV-2 infection in lung, intestine, and kidney epithelial cells, from proteolytic activation to membrane fusion. In agreement with other reports (7, 25), our results showed that SARS-CoV-2 infection was sensitive to inhibitors of TMPRSS2 and cathepsin L. We further found that blocking TMPRSS2 abrogated infection even when the cells were expressing cathepsin L, indicating that the virus does not reach endolysosomal cathepsins when TMPRSS2 is present. Others have shown that infection by MERS pseudo-viruses was suppressed by trypsin-like protease inhibitors in the presence of the tetraspanin CD9, while entry was unaffected but rather blocked by cathepsin inhibitors in the absence of CD9 (34). These authors proposed that tetraspanins condense CoV entry factors into localized positions on or close to the cell surface, allowing rapid and efficient activation of viral fusion (35).

We observed that SARS-CoV-2 used two distinct routes to enter cells, one fast (~10 min) which corresponded to the timing of TMPRSS2 activation, and the second slower (40-50 min) corresponding to cathepsin L priming. Although other cellular factors are likely necessary, our results support the view that TMPRSS2 is a major determinant of the SARS-CoV-2 fast entry track. Similar observations have been made for the human CoV 229E (hCoV-229E), which prefers cell-surface TMPRSS2 to endosomal cathepsins for cell entry (36–38).

It is clear from our data that, in the presence of TMPRSS2, SARS-CoV-2 did not rely on endosomal acidification for infectious penetration. Concanamycin B, which specifically inhibits vATPases and elevates endosomal pH, affected UUKV, an enveloped virus that penetrates host cells by acid-activated membrane fusion (26), but not SARS-CoV-2. This was consistent with reports that TMPRSS2 processes CoV S and other substrates at or nearby the plasma membrane (39, 40), i.e., at neutral pH. Using aprotinin, we found that half of the bound viral particles required 5-10 min to pass the TMPRSS2-dependent step. We cannot completely exclude that aprotinin was not instantaneously effective when it was added to the infected cells. In this case, the timing of TMPRSS2-requiring step was therefore faster. SARS-CoV-2 activation and penetration would then likely take place at the plasma membrane following proteolytic activation, as proposed for hCoV-229E and MERS-CoV (37, 41).

An alternative scenario would be that SARS-CoV-2 is sorted into the endocytic machinery regardless of the TMPRSS2 expression. The time course of TMPRSS2-requiring step resembled that of cargo sorted into EEs, *circa* 5-10 min (31). Another observation supporting this hypothesis was that colcemid hampered infection. This drug perturbates LE maturation by disrupting the microtubule network, and in turn, causes the accumulation and dysfunction of EEs (26). Such a strategy has been proposed for reoviruses, which use similar uptake but different trafficking depending on whether viral particles are activated or not (42). Like other CoVs (39), more functional investigations are required to determine, where exactly, from the plasma membrane or EEs, SARS-CoV-2 enters the cytosol of TMRPSS2+ cells, and whether the processing of the S protein is followed by transport of the virus to downstream organelles for penetration.

In the absence of TMPRSS2, it was evident that SARS-CoV-2 was dependent on endocytosis and transport through the late endosomal system for infectious penetration. Infectious entry was inhibited by endosomal-pH neutralizing drugs. Impairing LE maturation by either colcemid or the expression of Rab7a T22N affected SARS-CoV-2 infection. The sensitivity to MG-132 mirrored observations with UUKV, IAV, and murine CoVs, which accumulated in cytosolic vesicles and failed to infect (26, 30, 43). Others have reported that SARS-CoV-2 depends on PIKfyve for the infection of 293T cells, a line devoid of TMPRSS2 (10). PIKfyve is a phosphoinositide kinase involved in the first stages of LE maturation. Collectively, our results indicate that SARS-CoV-2, like other CoVs (41, 44, 45), has a dependence on functional endolysosomes and cathepsins for infectious penetration when the viral particles are not activated at or near the cell surface.

Our results suggested that the proteolytic activation of the spike S protein was sufficient and necessary for SARS-CoV-2 fusion. The Vero cells used in our virus-mediated cell-cell fusion assay did not express TMPRSS2 on the cell surface. In this assay, exogenous furin failed to promote the syncytia formation, indicating that either furin was inefficient or not sufficient to achieve the full activation of the SARS-CoV-2 protein S at the plasma membrane. The S1/S2 site exhibits a RRAR motif instead the typical RX(R/K)R furin one, and a recent structural study support the view that the cleavage by furin at this site in the S trimers is rather low, about 30% (17, 46, 47). However, we found that, unlike furin, trypsin prompted the formation of syncytia, which rather supports the involvement of proteases in the target cells, such as TMPRSS2 and cathepsin L, to complete the proteolytic processing of the S protein. Others have shown that SARS-CoV-2, and also MERS-CoV, mediate cell-cell fusion at neutral pH without any further proteolytic treatment when target cells express TMPRSS2 (16, 48).

It is also apparent from our results that the SARS-CoV-2 progeny was not fully processed and activated. Trypsin pretreatment increased the virus infectivity. More work is evidently required to decipher the SARS-CoV-2 fusion mechanism. The list of the involved host cell proteases is most likely not restricted to TMPRSS2 and cathepsin L, as suggested by a recent biochemistry study (49). The S proteolytic activation might involve the cleavage of other sites than S1/S2 and S2’, similarly to what was found for the MERS-CoV protein S (40). It is, however, tempting to postulate that the cleavage between S1 and S2 is not complete on SARS-CoV-2 particles, with only one or two of the three S1/S2 sites cut by furin within S trimers. In this model, cutting all the S1/S2 sites would be achieved by proteases in target cells such as TMPRSS2 and cathepsin L. The fusogenic conformational change would then occur and be completed by the cleavage of the S2’ sites, therefore, unmasking the fusogenic units. The S1/S2 site significantly differs in amino-acid residues through CoVs (17) and highly likely influences the overall viral fusion process.

We found that the level of virus mRNA and infectious viral progeny released in the outer media was lower in the absence of TMPRSS2. The TMPRSS2-dependent entry mechanisms occurred faster than the cathepsin L-activated pathway, and it might be that the early route results in a more productive infection than the late-penetrating process. Separate studies support, at least for some CoV strains including HCoV-229E, the view that early entry results in productive infection, while late penetration would be an alternative, backup route (35, 37, 38). Other works on therapeutics have linked host cell proteases to CoV spread. Inhibitors of TMPRSS2, but not of cathepsins, effectively prevent the pathogenesis of SARS-CoV in mice, suggesting that SARS-CoV mainly uses cell surface proteases rather than endosomal cathepsins *in vivo* (50). The identification of all host cell proteases involved in SARS-CoV-2 and other CoV infection, as well as the tissues and organs that express them, remains an important objective to better understand viral propagation and induced diseases.

Intriguingly, SARS-CoV-2 showed a strong resistance to acidic buffers. Exposure to pH ~5.0 only marginally inactivated the virus, and infectivity was even rescued and enhanced by proteolytic treatment. In addition, trypsin activation appeared to protect the virus from acid inactivation, which could explain how it is found to infect the gastrointestinal tract *in vivo*. SARS-CoV-2 has evidently developed a remarkable ability to adapt to an acidic environment. Interestingly, low pH has been shown to switch the positioning of the receptor-binding domain in the SARS-CoV-2 S trimers, which could help the virus to escape the immune system (51). Overall, this property certainly confers the virus the ability to sustain a high infectivity, not only within endosomes to enter host cells, but also in the extracellular space, especially during the virus spread throughout the host.

Reports on the cell entry of SARS-CoV-2 and other CoVs often describe only one cell model system, and the literature in this field remains confusing in general. Our study recapitulates within a single investigation the SARS-CoV-2 entry process and provides an overview of the cellular mechanisms used by SARS-CoV-2 to penetrate and infect target cells. Although it remains to be confirmed under physiological conditions, we propose that SARS-CoV-2 can enter cells through two distinct, mutually exclusive pathways. When target cells express TMPRSS2, the virus is activated at or close to the cell surface and penetrates early in a pH-independent manner. When target cells are devoid of TMPRSS2, SARS-CoV-2 is endocytosed and sorted into the endolysosomes from where the virus is activated in a pH-dependent manner and penetrates the cytosol late. With the ability to subvert diverse cell entry routes, SARS-CoV-2 has likely found a way to expand the number of target tissues and organs, which certainly contributes to the broad tropism of the virus *in vivo*.

## Materials and Methods

### Cells

The African green monkey Vero kidney epithelial cells (ATCC CRL 1586), the human Caco-2 colorectal adenocarcinoma (ATCC HTB-37), the human Calu-3 lung adenocarcinoma (ATCC HTB-55), and the human epithelial lung cells A549 stably expressing ACE2 (A549*; a kind gift from Prof. Ralf Bartenschlager) were all maintained in Dulbecco’s modified Eagle’s medium (DMEM) supplemented with 10% fetal bovine serum (FBS) and 100 units.mL^−1^ penicillin, and 100 μg.mL^−1^ streptomycin. Baby hamster kidney cells (BHK-21) were grown in Glasgow’s minimal essential medium containing 10% tryptose phosphate broth, 5% FBS, 100 units.mL^−1^ penicillin, and 100 μg.mL^−1^ streptomycin. All cell lines were grown in an atmosphere of 5% CO_2_ in air at 37°C. All products used for cell culture were obtained from Thermo Fischer Scientific and Sigma-Aldrich.

### Viruses

SARS-CoV-2 (strain BavPat1) was obtained from Prof. Christian Drosten at the Charité in Berlin, Germany, and provided via the European Virology Archive. The virus was amplified in Vero cells and working stocks were used after three passages. Uukuniemi (UUKV) and Semliki forest (SFV) viruses were previously described and amplified in BHK-21 cells (52, 53). The MOI is given according to the titer determined by plaque- or foci-forming unit assay for each cell line. When indicated, the titer was obtained by TCID50.

### Antibodies, reagents, and plasmids

The mouse mAb against the SARS-CoV nucleoprotein NP (40143-MM05) was purchased from Sino biologicals and used at dilutions of 1:500 for flow cytometry analysis and 1:1,000 for titration in TCID50 assays. The rabbit polyclonal antibody U2 targets all the UUKV structural proteins and was used at a dilution of 1:4,000 for immunohistochemistry (54). The mouse mAb 8B11A3 against the UUKV nucleoprotein N was a kind gift from Ludwig Institute for Cancer Research (Stockholm, Sweden) (55). The mouse mAb against the SFV glycoprotein E2 was kindly provided by Prof. Margaret Kielian (Albert Einstein College of Medicine, USA). mAb 8B11A3 and mAb against SFV E2 were used at a dilution of 1:400 for flow cytometry analysis. The rabbit antibodies against TMPRSS2 (ab92323) and actin (A2066) were obtained from Abcam and Sigma, respectively. The mouse mAb against cathepsin L (BMS1032) and α-tubulin (T5158) were bought from Thermo Fisher Scientific and Sigma, respectively. Anti-mouse secondary antibodies were conjugated to Alexa Fluor (AF) 405 (Molecular Probes), AF488 (Molecular Probes), IRDye 700 (LI-COR), IRDye 800CW (LI-COR), and horseradish peroxidase (HRP; Vector Laboratories). Anti-rabbit secondary antibodies conjugated to IRDye 800CW were purchased from LI-COR. NH_4_Cl (Sigma), aprotinin (Cayman Chemical), and chloroquine diphosphate (Sigma) stocks were dissolved in water. Bafilomycin A1 (BioViotica), Concanamycin B (BioViotica), SB412515 (Cayman Chemical), colcemid (Cayman Chemical), and MG-132 (Selleck Chemicals) were all dissolved in DMSO. Furin and Trypsin were purchased from R&D and Sigma, respectively. Plasmids encoding EGFP-tagged Rab7a, Rab7a T22N, and Rab7a Q67L have been described elsewhere (26).

### Protein analysis

Cells were lysed with phosphate buffer saline (PBS) containing 0.1% Triton X-100 (Merck Millipore), according to a standard procedure (54). Cell lysates were then diluted in LDS sample buffer (Thermo Fisher Scientific) and analyzed by SDS-PAGE (Nu-PAGE Novex 10% Bis-Tris gels; Thermo Fisher Scientific). Proteins were subsequently transferred to polyvinylidene difluoride membranes (iBlot transfer stacks; Thermo Fisher Scientific). The membranes were first blocked with Intercept blocking buffer (LI-COR) and then incubated with primary antibodies against TMPRSS2, cathepsin L, actin, and α-tubulin, all diluted in Tris-buffered saline containing 0.1% Tween and Intercept blocking buffer (1:1,000, 1:400, 1:5,000, and 1:2,000, respectively). After extensive washing, the membranes were incubated with the corresponding secondary anti-species conjugated to either IRDye 700 and 800CW (both at 1:10,000) or HRP (1:1,000). Proteins were analyzed with a LI-COR Odyssey CLx scanner, or alternatively, detected with SuperSignal West Pico PLUS chemiluminescent substrate (Thermo Fisher Scientific) and an iNTAS ECL Chemostar analyzer.

### Virus infection

Cells were exposed to viruses at the indicated MOIs in the presence of 2% FBS for 1 h at 37°C. Virus input was then replaced by complete culture medium, and infected cells were incubated for 8 h before fixation. For virus-mediated cell-cell fusion, Vero cells were infected for 24 h. Cells that transiently express EGFP-Rab7a and related mutants were infected 18 h post-transfection. For pH-inactivation, citric acid, 2-(N-morpholino)-ethanesulfonic acid (MES), and 4-(2-hydroxyethyl)-1-piperazineethanesulfonic acid (HEPES) were used as buffers at 100 mM as follow, pH < 5.5, 5.5 < pH < 6.5, and 6.5 < pH, respectively. Virus inputs were exposed to buffers at the indicated pH for 10 min at 37°C and then to buffers at neutral pH prior infection. For furin- or trypsin-activation, SARS-CoV-2 was pretreated with furin (1 μg.mL^−1^) or trypsin (100 μg.mL^−1^), respectively, for 15 min at 37°C and allowed to infect cells. For inhibition assays, cells were pretreated with drugs for 30 min at 37°C, apart from colcemid pretreatment that lasted 3 h on ice, and then exposed to viruses in the continuous presence of the inhibitors. For inhibitor add-in time courses, virus binding to cells was synchronized on ice for 90 min. Cells were then rapidly warmed to 37°C, and SB412515 (10 μM), aprotinin (30 μM), NH_4_Cl (at indicated concentrations), Concanamycin B (50 nM), and MG-132 (at indicated concentrations) were added at indicated times. Cells were subsequently incubated at 37°C and harvested 8 h after the warm shift. Infection was monitored by either flow cytometry, fluorescence microscopy, or qRT-PCR. When infection was analyzed by microscopy, cells were seeded on Lab-Tek or iBIDI glass bottom 8-well chamber slides.

### DNA transfection

As previously described (56), Vero cells were transfected with 750 ng of plasmids using Lipofectamine 2000 (Invitrogen) in 24-well-plates according to the manufacturer’s recommendations and washed 5 h later.

### Immunofluorescence microscopy

Fluorescence microscopy was extensively described in (57). Briefly, infected cells were rinsed with PBS, permeabilized with 0.5% Triton X-100 (Sigma) for 15 min at room temperature (RT), and stained with primary antibodies diluted in PBS for 1h at RT. Subsequently, cells were extensively washed and incubated with secondary antibodies in the presence of 4′,6-diamidino-2-phenylindole (DAPI, Molecular Probes) for 45 min at RT. Samples were imaged with an epifluorescence microscope Nikon Eclipse Ti-S (Nikon), whilst a Leica TCS SP8 confocal microscope was used to image syncytia.

### Flow cytometry

The flow cytometry-based infection assay has been described previously (53). Briefly, infected cells were fixed with 4% formaldehyde for 30 min at RT and permeabilized with 0.1% saponin (Serva). Cells were then exposed to primary antibody at RT for 1 h, washed, and subsequently incubated with secondary anti-mouse antibodies at RT for another 1 h. Infected cells were quantified with a FACSCelesta cytometer (Becton Dickinson) and FlowJo software (TreeStar).

### Virus RNA quantification

As previously reported (58), RNA was harvested from cells using the NuceloSpin RNA extraction kit (Machery-Nagel) according to manufacturer’s instructions. The cDNA was synthesized using iSCRIPT reverse transcriptase (BioRad) from 250 ng of total RNA as per supplier recommendations. q-PCR was performed using iTaq SYBR green (BioRad) following the manufacturer’s instructions for the SARS-CoV-2 genome using the forward primer, GCCTCTTCTCGTTCC, and the reverse primer, AGCAGCATCACCGCC. HPRT1 was used as a housekeeping gene using the forward primer, CCTGGCGTCGTGATTAGTGAT, and reverse primer, AGACGTTCAGTCCTGTCCATAA.

### Virus titration by TCID50 assay

Confluent monolayers of Vero and Caco-2 cells in 96-well plates were infected with 10-fold serial dilutions of SARS-CoV-2. Infected cells were fixed 24 hpi and subjected to immunostaining using the primary mouse mAb anti-SARS-CoV-2 NP and then the secondary anti-mouse antibody 800CW (1:10,000). Samples were finally scanned on LI-COR.

### Cell-cell fusion

Infected cells were washed in PBS and treated with DMEM containing 0.2% bovine serum albumin (Gibco) buffered at pH 7.4, 6.0, or 5.0 using 30 mM of HEPES, MES, or citric acid, respectively, for 5 min at 37°C. Alternatively, infected cells were exposed to furin (1 μg.mL^−1^) and trypsin (100 μg.mL^−1^) for 5 min at 37°C, and when indicated, followed by acidification of the culture medium as described above. Subsequently, cells were washed and incubated in complete medium for 50 min, and the cytosol stained with CellMask Deep Red (1:1,000, Molecular Probes) for 10 min at 37°C. After fixation, cells were rinsed with PBS, and nuclei stained with Hoechst 33258 (0.5 μg.mL^−1^, Thermo Fisher Scientific). Syncytia were monitored by fluorescence microscopy as described below. Fusion was quantified by counting the number of cells and nuclei present in a microscope field. A fusion index (*f*) was calculated according to the equation *f* = (1 – [*c*/*n*]), where *c* is the number of cells in a field after fusion and *n* the number of nuclei. An average field contained 50-60 nuclei.

### Statistical analysis

The data shown are representative of at least three independent experiments. Values are given as the means of duplicate ± standard error of mean or triplicates ± standard deviations. Graph plotting of numerical values, as well as the statistics, were achieved with GraphPad Prism v5.00 (GraphPad Software). Statistical methods and parameters are indicated in the figure legends when applicable. P-values are shown when statistical differences are significant.

## Acknowledgments

This work was supported by grants from CellNetworks Research Group funds, Heidelberg, and from the Deutsche Forschungsgemeinschaft (DFG) project numbers LO-2338/1-1 and LO-2338/3-1) to PYL, project numbers 415089553 (Heisenberg program), 240245660 (SFB1129), 278001972 (TRR186), and 272983813 (TRR179) to SB, and project 416072091 to MS. This work was also supported by INRAE starter funds, IDEX-Impulsion 2020 (University of Lyon), and FINOVI (Fondation pour l’Université de Lyon), all to PYL. We acknowledge funding from the German Academic Exchange Service (DAAD, Research Grant 57440921) to PD. We thank Felix Rey and Ari Helenius for fruitful discussions.

